# Regulatory preconditioning for the evolution of C_4_ photosynthesis revealed by low CO_2_ treatment of Arabidopsis thaliana

**DOI:** 10.1101/2023.07.10.548402

**Authors:** Fenfen Miao, Noor UI Haq, Ming-Ju Amy Lyu, Xin-Guang Zhu

## Abstract

Low CO_2_ condition was considered a preconditioning or selection pressure for C_4_ evolution. However, it remains elucidated how low CO_2_ condition contribute to the evolutionary assembly of the C_4_ pathway. We conducted a systematic transcriptomics and metabolomics analysis under short-term low CO_2_ condition and found that *Arabidopsis* grown under this condition showed increased expression of most genes encoding C_4_- related enzymes and transporters. Low CO_2_ condition increased NH_4_^+^ content in leaves; as expected, photorespiratory and ammonia refixing pathways were enhanced. Furthermore, we found that compared to low CO_2_ condition, in vitro treatment with NH_4_^+^ induced a similar pattern of changes in C_4_ related genes and genes involved in ammonia refixation. This supports that increased expression of C_4_ genes induced by low CO_2_ condition can supply carbon skeleton for ammonia recycling. This study provides new insight into the regulatory preconditioning which may have facilitated the evolution of C_4_ photosynthesis under low atmospheric CO_2_ environments.

## Introduction

Although C_4_ plants only have 8000 species, but it contributes to 23% of the earth’s terrestrial primary productivity (Still et al., 2003; Sage et al., 2007). With over 60 distinct evolutionary origins (Sage et al., 2011; Sage et al., 2018), C_4_ photosynthesis is an excellent example of convergent evolution, which attracts attentions among biologists to study the evolutionary history of C_4_ photosynthesis and the reason underlying so many independent origins (Sage, 2001; Sage et al., 2011). According to the present models of C_4_ evolution (Sage, 2011; Sage, 2012), relocation of photorespiratory CO_2_ release to the bundle sheath cell (BSC) from the mesophyll cell (MC) is an important early step during the evolution of the C_4_ photosynthesis, which is also termed as C_2_ photosynthesis (Vogan et al., 2007; Sage, 2012). This mechanism is characterized by a photorespiratory CO_2_ pump which is realized by restricting glycine decarboxylation (GDC) to the BSC and glycine, a two-carbon compound, that originated from photorespiration to diffuse to the BSC for decarboxylating and concentrating CO_2_. CO_2_ is enriched threefold by this pathway (Keerberg et al., 2014). Immunogold experiments supported this hypothesis, which showed GDC is expressed exclusively in the BSC of some intermediates, such as *Moricandia arvensis* (Rawsthorne et al., 1988; Schluter et al., 2017) and *Heliotropium* (Vogan et al., 2007). Due to the enrichment of CO_2_, Ribulose-1,5-bisphosphate carboxylase/oxygenase (Rubisco) works more efficiently in the BSC leading to an overall increase in biomass production under condition that support higher rates of photorespiration (Heckmann et al., 2013; Mallmann et al., 2014), such as low CO_2_, drought and high temperature (Wingler et al., 1999; Carmo-Silva et al., 2008; Li et al., 2014; Sage et al., 2018).

Accompanied with the release of CO_2_ by GDC during photorespiration, ammonia (NH_4_^+^) is also released (Bauwe et al., 2010), which needs to be refixed since it is toxic (Krogmann et al., 1959). Photorespiratory ammonia can be integrated into amino acids via the chloroplastic isoform Glutamine synthetase 2 (GS2) and ferredoxin-dependent GOGAT (Fd-GOGAT) (Lam et al., 1996). This process forms the photorespiratory ammonia cycle (Keys, 2006). Photorespiratory CO_2_ pump results in ammonia imbalance between MC and BSC, which needs to be solved through metabolic shuttles (Nagatani et al., 1971; Lancien et al., 2000; Liu and von Wiren, 2017). Mallman (Mallmann et al., 2014) showed that there are multiple solutions to the ammonia imbalance, with one option, an alanine-pyruvate shuttle, providing a bridge toward C_4_ photosynthesis metabolism. Therefore, ammonia refixation induced by high rate of photorespiration was proposed a driving force for C_4_ evolution. However, it raises a question regarding how these metabolic shuttles which recycle ammonia emerge during C_4_ evolution. Considering that these metabolic shuttles for ammonia recycling require a rather larger scale rewiring of the plant’s primary metabolism, quickly evolving such a mechanism when ammonia imbalance in C_2_ plants appeared would be rather difficult. Importantly, C_4_ photosynthesis mostly emerged around 30 million years ago and originated independently more than 60 times (Sage et al., 2011), there seems little likelihood for such complex metabolic pathways and its associated regulatory mechanisms to evolute multiple times in such a short time. Previous study suggested that all genes involved in C_4_ photosynthesis are present in C_3_ ancestral plants and play important house-keeping roles (Aubry et al., 2011). However, whether the NH_4_^+^ recycling pre-existed or whether it plays crucial roles before C_4_ evolution remains to be tested.

The 30 million-year spread in the timing of many C_4_ plant origins following the Oligocene CO_2_ decline indicates that low CO_2_ condition maybe a preconditioning for C_4_ evolution (Ehleringer et al., 1991; Ehleringer et al., 1997; Christin et al., 2008). Low CO_2_ condition enhances photorespiration (Li et al., 2014). However, there is no evidence suggesting how this low CO_2_ preconditioning contributes to the evolutionary formation of the C_4_ pathway. In this study, we attempt to answer this question through combined transcriptomics and metabolomics analysis. One caveat in this study is that we only test whether the regulatory mechanisms on gene expression pre-existed before C_4_ evolution, while the pre-existing sequence variations on key enzymes of C_4_ photosynthesis, such as PEPC (Westhoff and Gowik, 2004; Schluter and Weber, 2020) were not studied, given that under short-term low CO_2_ treatment, changes in enzyme properties would not be expected. The results suggest low CO_2_ condition can effectively induce metabolic rewiring which by providing carbon skeletons and creates an effective ammonia recycling pathway in the C_3_ model plant *Arabidopsis*. We also found that ammonia accumulation and refixation plays an important role during this metabolic rewiring. This ability to rewire plant primary metabolism can serve as regulatory preconditioning underlying multiple parallel evolutions of C_4_ photosynthesis.

## Materials and Methods

### Plant material and growth conditions

*Arabidopsis* used in this study is the Columbia (Col-0) ecotype. Seeds were surface-sterilized by incubation with 10 % (v/v) bleach diluted with ethanol, washed three times with sterile distilled water, and then sown in plastic Petri dishes (diameter 90 mm, depth 20 mm) containing 1/2 MS medium, solidified with 0.7 % (w/v) agar and supplemented with 1.5 % (w/v) sucrose (pH 5.8). The seeds were placed in the dark at 4 °C for three days and germinated for six days in incubators with a photosynthetic photon flux density (PPFD) of 100 μmol m^-2^s^-1^ (10 h light/14 h dark cycle) at 22 °C. After this, seedlings were transferred to *Pindstrup* soil and grown in a Percival incubator (Nihonika, Japan) in which CO_2_ gas was accurately and stably controlled.

### Low CO_2_ treatment

The CO_2_ condition of 100 ppm (low CO_2_, LC) and 400 ppm (normal CO_2_, NC) were applied in two separate Percival incubators (Nihonika, Japan) and maintained throughout this study. *Arabidopsis* were grown for 14 days under a PPFD of 100 μmol m^-2^s^-1^ (10 h light/14 h dark cycle) at 22 °C and relative humidity of 65 % (Li et al., 2014). After, half of the plants were transferred to low CO_2_ condition for six days. New fully expanded leaves under low CO_2_ and normal CO_2_ treatment were sampled for transcriptomic and metabolomics analysis as well as the measurements of chlorophyll content and free ammonia contents.

### Ammonia treatment

To study the effect of ammonia ions on gene expression, seeds were surface-sterilized and sown in a control check (CK) plastic Petri dish (diameter 90 mm, depth 20 mm) in a growth chamber for 10 days at a PPFD of 100 μmol m^-2^s^-1^ (10 h light/14 h dark cycle) at 22 °C. The composition of the CK was described in (Li et al., 2013), which is composed of 2 mM KH_2_PO_4_, 5 mM NaNO_3_, 2 mM MgSO_4_, 1 mM CaCl_2_, 0.1 mM Fe-EDTA, 50 μM H_3_BO_3_, 12 μM MnSO_4_, 1 μM ZnCl_2_, 1 μM CuSO_4_, 0.2 μM Na_2_MoO_4_, 0.5 gl^-1^ MES, 1% sucrose, and 0.8% agarose (pH 5.7, adjusted with 1 M NaOH). Half of the seedlings were transferred to the new CK petri dish and the remaining seedlings were transferred to new CK plates which were additionally supplied with 15 mM or 30 mM (NH_4_)_2_SO_4_. Nine seedlings were put on one Petri dish and considered as one biological replicate during sampling. After three days of treatments, leaves were harvested and immediately put into liquid nitrogen and stored at –80 °C until RNA extraction. We also put seedlings on the CK plates which were additionally supplied with 30 mM or 60 mM NH_4_CL, 15 mM or 30 mM K_2_SO_4_, 30 mM or 60 mM KNO_3_, and 120 mM mannitol to test the effect of other ions (Li et al., 2019).

### Measurement of free ammonia content in a leaf

Ammonia content was determined with the colorimetric method based on the phenol hypochlorite assay (SolÓRzano, 1969; Sarasketa et al., 2014). Briefly, leaves were collected, weighed, and snap-frozen in liquid nitrogen, and then ground with 5 mm zirconia beads. 1 ml H_2_O was added to the frozen powder. The mixtures were incubated at 80 ℃ for 10 min and then centrifuged at 4000 *g* and 4 °C for 20 min. 100 µl of supernatant was mixed with 200 μl of 0.33 M sodium phenolate, 100 µl of 0.02% sodium nitroprusside; then 200 µl of 2% sodium hypochlorite were added to the mixture. The mixture was incubated at room temperature for 30 min followed by reading absorbance at a wavelength of 630 nm in a spectrophotometer.

### Gene expression analysis

For gene expression profiling, we took illuminated new fully expanded leaf samples. Four biological replicates were harvested and frozen in liquid nitrogen immediately and stored at –80 °C before RNA extraction. For RNA extraction, the PureLink RNA Mini Kit (Life Technologies Corporation, USA) was used to extract RNA following the manufacturer’s protocol, which includes the removal of gDNA from RNA. The quality of purified RNA was assessed using an Agilent 2100 Bioanalyzer (Agilent, USA) and the RNA samples with RNA Integrity Number (RIN) higher than 7 were used for RNA library construction. RNA libraries were prepared based on standard Illumina (Illumina, Inc., USA) protocols and sequenced using the Illumina X Ten platform in paired-end 150 bp mode. The quality of RNA-seq data (fastq files) was assessed by the FastQC software (https://www.bioinformatics.babraham.ac.uk/projects/fastqc/). RNA-seq analysis was performed by the STAR software (Dobin et al., 2013) with the *Arabidopsis* reference genome as well as a gene transfer format (GTF) file (downloaded from EnsemblPlants http://plants.ensembl.org/). After generation of the genome index, RNA-seq reads were aligned by STAR with the ‘--quantMode GeneCounts’ option to count reads per gene. Differentially expressed genes (DEGs) were determined by the R package ‘DESeq2’ (Love et al., 2014) which use the Benjamini-Hochberg (BH) adjustment for multiple testing problem with the read counts reported by STAR (Dobin et al., 2013). DEGs between treatment and control were analyzed. Only genes with the adjusted *P*-value <0.05 (Padj) were considered as DEGs in each treatment.

### Functional analysis of differentially expressed genes

Gene transcript abundances and metabolite abundances were visualized applying R package ‘heatmap’ (https://cran.r-project.org/web/packages/pheatmap/index.html). Permutation of Pearson correlations was conducted used the R package Permutation test. Volcano plot and Venn diagrams were plotted applying online tool Bioinformatics (https://www.bioinformatics.com.cn). To identify functional categories of differentially expressed genes, Gene Oncology (GO) and Kyoto Encyclopedia of Genes and Genomes (KEGG) pathway enrichment analyses were performed using the Database for Annotation, Visualization and Integrated Discovery (DAVID) (https://david.ncifcrf.gov/). Data were analyzed using the two-tailed Student’s *t-test*. The term significant is used here for differences or correlations confirmed at *\* P* < 0.05 or better. Details are provided in the figure legends.

### Liquid Chromatography/Mass Spectrometry and Metabolomics analysis

To enable the sampling of leaves grown under low CO_2_, we modified the glass door of a Percival incubator to ensure the stability of CO_2_ condition during leaf sampling. Specifically, we cut two spherical openings in the middle of the glass door and installed two plastic gloves used for sampling.

The Liquid Chromatography/Mass Spectrometry (LC-MS/MS) experiments were performed following (Wang et al., 2014; Arrivault et al., 2019). Fully expanded leaves under light were sampled. For each sample, a leaf area of 1.7 cm^2^ was sampled and frozen in liquid nitrogen instantaneously. The same leaf position was sampled to measure the fresh weight to calculate specific leaf weight. All leaf samples were cut *in situ* and immediately transferred into a pre-frozen 2 mL EP tube, then stored in liquid nitrogen for metabolite extraction. After grinding, each sample was fully dissolved with 800 μL extraction buffer (methanol: chloroform = 7:3 (v/v), -20 ℃ pre-cooling) and incubated under -20 ℃ for 3 hours. Then 560 μL distilled water (ddH_2_O) was added and mixed with each sample, and 800 μL supernatant was extracted after centrifugation (×2200g, 10min, 4℃). After that, 800 μl buffer (methanol: ddH_2_O = 1: 1(v/v), 4 ℃ pre-cooling) was mixed with the sample for another extraction. For each sample, 1.6 mL supernatant in total was obtained by filtering the extraction buffer with a 0.2 μM nylon filter. Among them, 1 mL was used for MS/MS analysis, and 20 μL was used for the QC sample. All extraction operations were performed on ice.

Luna NH_2_ column (3μm, 100mm*2mm, Phenomenex co. Ltd, USA) was used in the liquid chromatography. The Liquid Chromatography gradient was set with eluent A, which has 10 mM Ammonia acetate and 5% (v/v) acetonitrile solution, with the pH adjusted to 9.5 using ammonia water and eluent B (acetonitrile): 0-1 min, 15% A; 1-8 min, 15-70% A; 8-20 min, 70-95% A; 20-22 min, 95% A; 22-25 min, 15% A. During the mass spectrometry analysis, QTRAP 6500+ (AB Sciex, co. Ltd, USA) was used in the MRM model with all parameters used following (Wang et al., 2014; Arrivault et al., 2019). The condition of all metabolites in samples were calculated based on the “condition-peak area” curve of standard samples and converted to nmol·gFW-1 with specific leaf weight (Arrivault et al., 2019).

### Real-time RT–qPCR

The RT–qPCR analysis was conducted as described previously (Chen et al., 2020). The RNA was extracted with the same procedure as described in Gene expression analysis. For reverse transcription, 0.2 μg RNA was reverse transcribed with TranScript One-Step gDNA Removal and cDNA Synthesis SuperMix (TransGen Biotech, China), then SYBR Green Real-Time PCR Master Mix (Yeasen, China) was used for qPCR on the CFX 96 system (Bio-Rad) following the manufacturer’s protocol. The ACTIN 8 (AT1G49240) gene was used as a reference for mRNA normalization. The comparative cycle threshold (Ct) method was used to evaluate the relative gene expression levels. The primers used for the expression analysis are listed in Table S1.

### Measurement of Chlorophyll Content

Leaf discs (0.8 cm^2^) were taken from fully expanded leaves, and chlorophyll was extracted with 80% acetone at 4 °C for 24 h in darkness, then the supernatant was used for the absorbance measurement at 652 nm in a spectrophotometer (Arnon, 1949). The total chlorophyll content was calculated with the following formula:

Total chlorophyll (μg/cm^2^) =34.5× OD652 (μg/ml) ×V (ml) / leaf area (cm^2^)

## Results

### The genes related in primary metabolism were induced under low CO_2_ condition

To study the effect of low CO_2_ condition on C_3_ plants, we examined the physiological, transcriptomic, and metabolomic changes of *Arabidopsis* under two CO_2_ conditions: low CO_2_ with 100 ppm (LC) and normal CO_2_ with 400 ppm (NC). Plants grown under low CO_2_ condition had lower biomass and decreased chlorophyll than those grown under normal CO_2_ (Fig. S1A, B).

The differentially expressed genes (DEGs) were identified between low CO_2_ and normal CO_2_ treatments (Dataset S1). 2976 genes were upregulated and 2892 genes were downregulated (Fig. S2A). Gene Ontology (GO) enrichment analysis shows that hexose catabolic process, sugar-phosphatase activity, carbohydrate phosphatase activity, amyloplast, carotene biosynthetic and abscisic acid metabolic process were significantly enriched in upregulated genes (Fig. S2B, Dataset S2), whereas sulfur compound and jasmonic acid-mediated signaling pathway were enriched in the downregulated genes (Dataset S2).

In addition, the significantly upregulated genes were mainly participated in metabolic process, including biosynthesis of secondary metabolites, glycolysis / gluconeogenesis, starch and sucrose and carbon metabolism by KEGG analysis (Fig. S3C Dataset S2). These results show that low CO_2_ induced changes to gene expression involved in primary metabolism.

### Most C_4_ related genes are upregulated under low CO_2_ condition in *Arabidopsis*

Notably, under low CO_2_, 23 of 26 genes associated with C_4_ metabolism were significantly increased (Fig. 1; Dataset S1), including cytoplasmic carbonic anhydrase (CA2, AT5G14740; CA4, AT1G70410) and chloroplast-localized carbonic anhydrase (CA5, AT4G33580). The gene expression of phosphoenolpyruvate carboxylase (PEPC, At2G42600, AT1G53310), PEPC kinase (PPCK1, AT1G08650), chloroplast/mitochondrial NAD-dependent malate dehydrogenase (pNAD-MDH, AT3G47520; mMDH1, AT1G53240), pyruvate orthophosphate dikinase regulatory protein (PPDK-RP, AT4G21210), alanine aminotransferase (AlaAT1, AT1G17290), and aspartate aminotransferase (AspAT, AT4G31990, AT5G19550, AT2G22250) were also dramatically upregulated (Fig. 1; Dataset S1). In addition, the genes encoding C_4_ related transporters were upregulated significantly, which includes phosphoenolpyruvate/phosphate translocator (PPT1, AT3G01550; PPT2, AT5G33320), dicarboxylate carriers (DIC, AT4G24570; DIC2, AT2G22500), inorganic pyrophosphatase 2 (PPA2, AT2G18230), plasma membrane protein (PIP1, AT3G61430), and mesophyll envelope protein (MEP2, AT5G23890; MEP3/4, AT5G12470), dicarboxylate transporters (DIT1, AT5G12860; DIT2.1, AT5G64290). Overall, our results show preconditioning of low CO_2_ condition contribute to the evolutionary assembly of the C_4_ pathway by induced mostly C_4_ related genes.

**Fig. 1.**
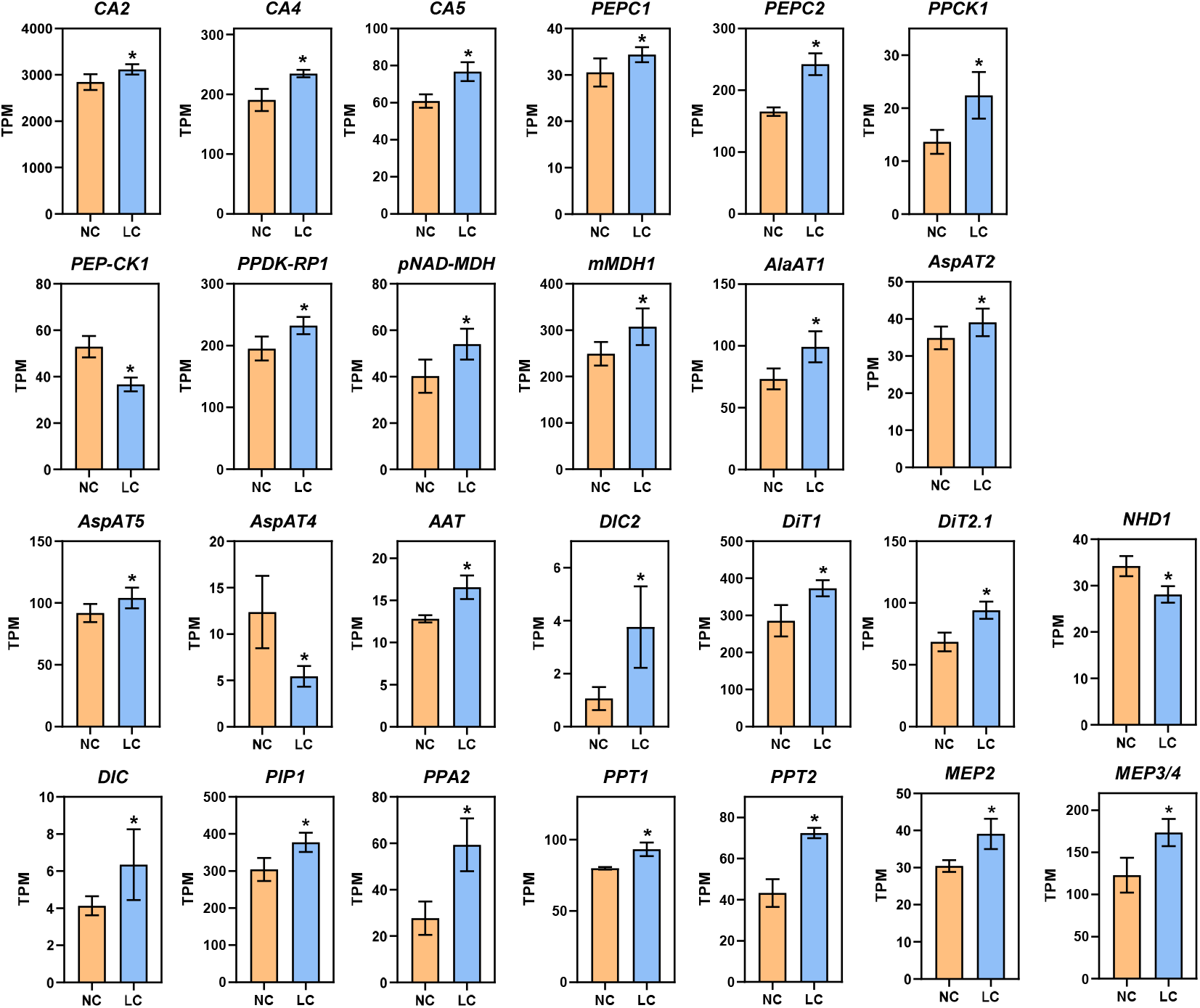
Low CO_2_ induction of C_4_ related genes in *Arabidopsis.* RNA-seq data are plotted as transcript per million (TPM). Gene expression is shown from samples collected from NC (normal CO_2_, 400 ppm) and LC (low CO_2_, 100 ppm). *CA2, Carbonic Anhydrase 2; CA4, Carbonic Anhydrase 4; CA5, Carbonic Anhydrase 5; PEPC1, Phosphoenolpyruvate Carboxylase 1; PEPC2, Phosphoenolpyruvate Carboxylase 2; PPCK1, PEPC Kinase1; PEP-CK1, Phosphoenolpyruvate Carboxykinase 1; RPDK- RP1, PPDK Regulatory Protein 1; pNAD-MDH, chloroplast NAD-Dependent Malate Dehydrogenase; mMDH1, mitochondrial Malate Dehydrogenase; AlaAT1, Alanine Aminotransferase1; AspAT4, Aspartate Aminotransferase 4; AspAT5, Aspartate Aminotransferase 5; AspAT2, Aspartate Aminotransferase 2; AAT, Aspartate Aminotransferase; PIP1, Plasma Membrane Protein; DIC2, Dicarboxylate Carriers 2; DiT2.1, Dicarboxylate Transport 2.1; DiT1, Dicarboxylate Transporter 1; DIC, Dicarboxylate Carriers; PPA2, Inorganic Pyrophosphatase 2; PPT1, Phosphoenolpyruvate Translocator 1; PPT2, Phosphoenolpyruvate Translocator 2; NHD1, Na+/H+ antiporter; MEP2, Mesophyll Envelope Protein 2; MEP3/4, Mesophyll Envelope Protein 3/4.* Data are shown as mean ± s.d (replications *n*=4). *\**, *adjusted P- value* <0.05 which was determined by DESeq2.

We validated 12 genes that are involved in primary metabolism through RT-qPCR. The results of the RT-qPCR were consistent with the RNA-seq analysis as shown in Figure S3A, with a Pearson correlation coefficient of 0.75. (Fig. S3B).

### Photorespiratory and ammonia refixation pathway are enhanced under low CO_2_ condition

Photorespiration was enhanced under low CO_2_ condition (Li et al., 2014). Photorespiratory ammonia refixation are highly integrated with photorespiratory metabolism (Keys et al., 1978). Therefore, we investigated the expression of genes related to photorespiration and ammonia refixation pathways. Almost all photorespiratory genes, including glycolate oxidase (GOX2, AT3G14415), glutamate: glyoxylate aminotransferase (GGT2, AT1G70580), alanine: glyoxylate aminotransferase (AGT1, AT2G13360), serine hydroxymethyl transferase (SHM2, AT5G26780; SHM4, AT4G13930), glycine decarboxylase P-protein (GLDP1, AT4G33010; GLDP2, AT2G26080), glycine decarboxylase L-protein (mtLPD2, AT3G16950; mtLPD4, AT4G16155), and mitochondrial transporter A BOUT DE SOUFFLE (Bou, AT5G46800) (Eisenhut et al., 2013), were significantly upregulated (Fig. 2A). The gene coding for 2- phosphoglycolate phosphatase (PGLP1, AT5G36700; PGLP2, AT5G47760) showed dramatically decrease (Fig. 2A). Concurrent metabolomics analysis shows that the shift to low CO_2_ condition led to markedly larger intracellular pools of 2-phosphoglycolate (2- PG), glycolate, glycine, and glycerate, which are intermediates in the photorespiratory pathway (Fig. 2A). The increased metabolite levels corroborate with the upregulation of photorespiratory gene expression. In addition, the NH_4_^+^ levels in leaves were increased by one-fold compared to normal CO_2_ treatment (Fig. 2B).

**Fig. 2.**
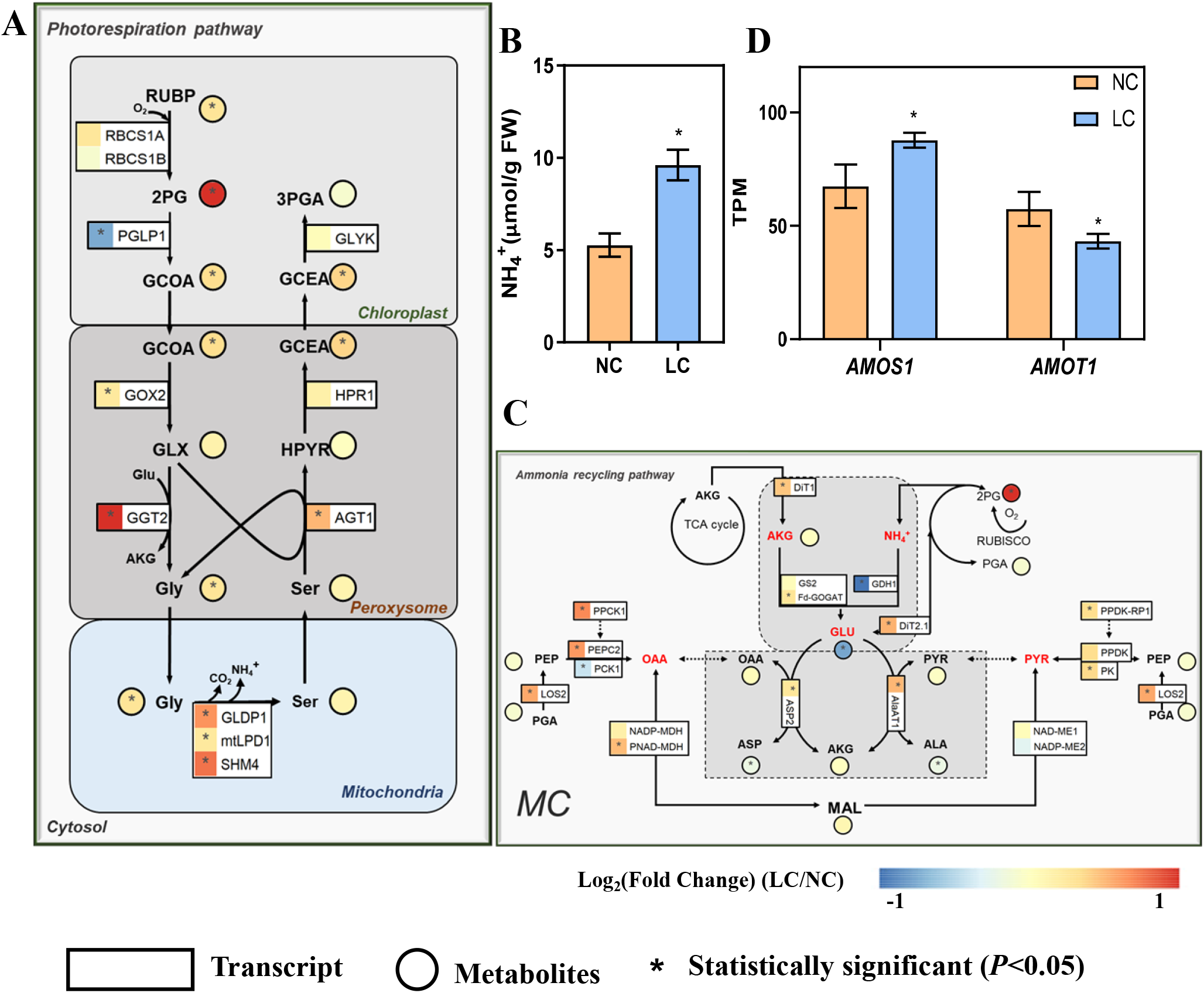
Changes of the gene expression and metabolites concentration on photorespiratory and ammonia recycling pathways under low CO_2_. (A) Photorespiratory pathway. (B) NH_4_^+^ content in the leaf (replications *n*=4). (C) Ammonia recycling pathway. (D) The changes of expression of ammonia-sensitive genes. AMOS1 (AT5G35220,*Ammonia- overly-sensitive 1*), *AMOT1 (*AT3G20770,*Ammonia tolerance 1).* The relative changes of gene expression and metabolites are shown in log_2_Fold change in heatmap, which use the same heatmap scale. For gene expression (A, C, D) differentially expressed genes (DEGs) were determined by the R package ‘DESeq2’ with the *adjusted P-value* <0.05 (replications *n*=4). For metabolites (A, C) and free NH_4_^+^ content in the leaf (B) which were determined by two-sided Student’s *t-test* (replications *n*=5) and * indicates *P*<0.05. Data are shown as mean ± s.d (B, D). The symbols of genes in these metabolic pathways are from the Arabidopsis Information Resource (TAIR) Database (http://www.arabidopsis.org/), and also can get from Dataset S1.

For ammonia refixation pathway, the gene coding Fd-GOGAT (AT5G04140), which is involved in ammonia refixation, also showed significantly higher expression level under low CO_2_ condition (Fig. 2C). DIT1 and DIT2.1, which work in concert in photorespiratory carbon/nitrogen metabolism; the lack of either DiT2.1 or DiT1 led to growth impairment under photorespiratory condition (Kinoshita et al., 2011), were increased markedly (Fig. 2C). Moreover, the gene expression of AspAT and AlaAT, which are involved in the transamination, showed significantly increase (Fig. 2C). The concentration of 2-oxoglutarate (2-OG) showed no significant change (Fig. 2C), whereas the concentration of glutamate was reduced by half (Fig. 2C). The concentration of oxaloacetate (OAA) and pyruvate were not changed under low CO_2_ condition, while those of aspartate and alanine were slightly decreased (Fig. 2C).

Under low CO_2_ condition, the expression of the gene *AMOS1* (*Ammonia-overly sensitive 1*) (Li et al., 2012), encoding a plastid metalloprotease that confers sensitivity to ammonia, significantly increased as depicted in Figure 2D. Conversely, the expression of AMOT1 *(Ammonia tolerance 1*) (Li et al., 2019), which is crucial for mitigating the effects of ammonia toxicity and confers greater tolerance to NH_4_^+^ in mutants than the wild type, was markedly reduced (Fig. 2D). Overall, the expressions of photorespiratory, ammonia recycling pathway and ammonia-sensitivity genes were induced, indicating *Arabidopsis* was subjected to ammonia stress under low CO_2_ condition.

### Low CO_2_ condition reprogramming primary carbon metabolism

The KEGG enrichment analysis of DEGs showed that low CO_2_ mainly affects genes related to primary carbon metabolism (Dataset S2). Among the affected genes, a vast majority of those involved in the tricarboxylic acid cycle (TCA) were markedly upregulated, specifically aconitase (ACO1, AT4G35830), isocitrate dehydrogenase 1 (IDH1, AT4G35260), 2-oxoglutarate dehydrogenase (OGDC, AT3G55410), succinate dehydrogenase (SDH, AT5G66760), and mMDH1 (Fig. 3A, Dataset S1). However, we observed a significant reduction of the succinate and fumarate in comparison to normal CO_2_ condition (Fig. 3A).

**Fig. 3.**
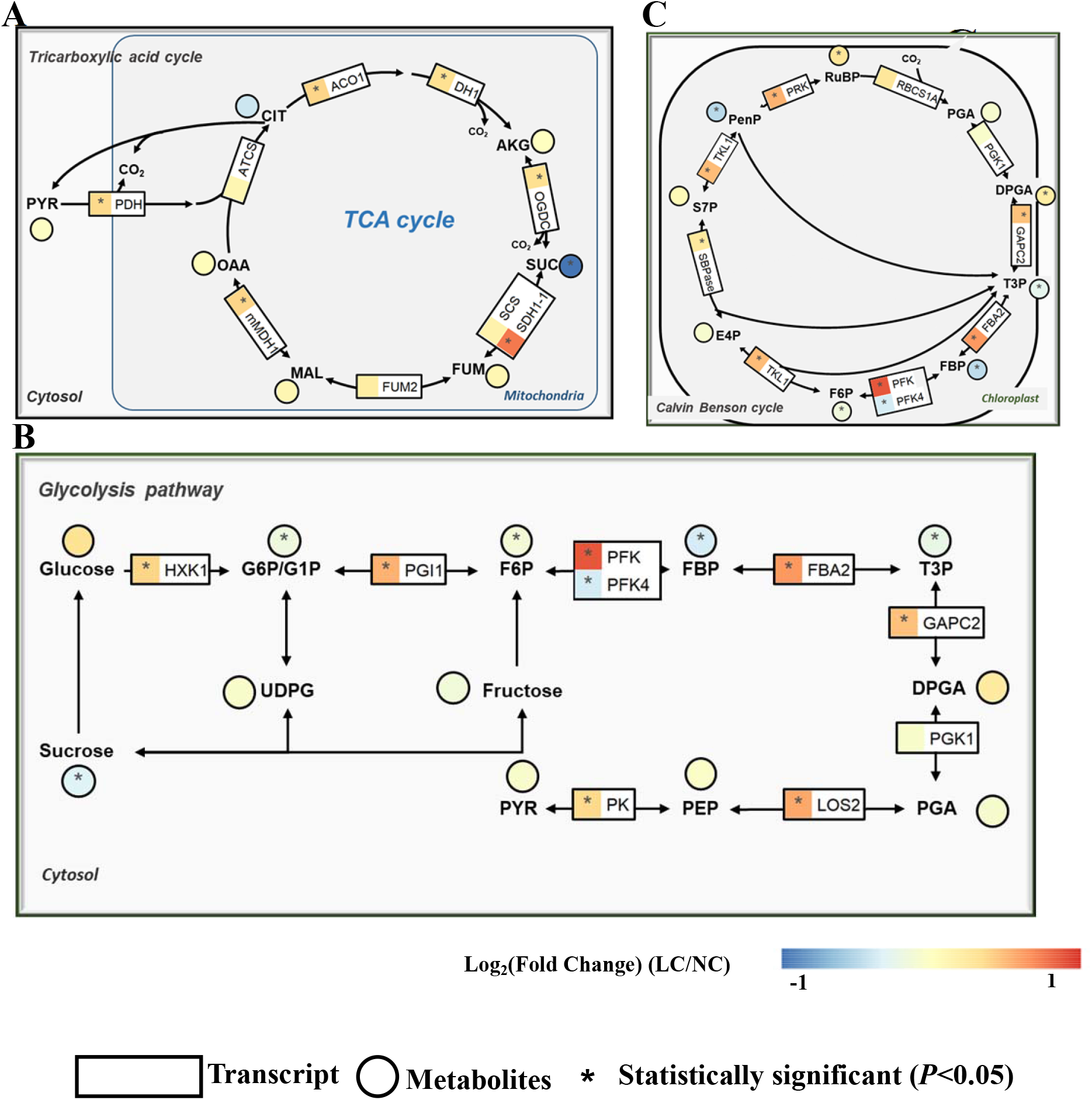
The effect of low CO_2_ on gene expression and metabolites condition in carbon metabolism pathways. (A) Tricarboxylic acid cycle (TCA cycle). (B) Glycolysis pathway. (C) Calvin-Benson-Bassham (CBB) cycle. The details are same as Fig. 2.

The expression levels of genes associated with the glycolysis pathway showed similar trends of changes as those observed among the TCA cycle genes. Notably, most of these genes demonstrated significant upregulation while metabolites like sucrose, glucose-6 phosphate (G6P), fructose-6-phosphate (F6P), fructose-1:6-Bisphosphate (FBP), and triose phosphate (T3P) were significantly reduced under low CO_2_ treatment (Fig. 3B). We observed increased expression levels of enolase (LOS2, AT2G36530), pyruvate kinase family protein (AT5G08570; AT2G36580), and pyruvate dehydrogenase (PDH-E1 BETA, AT1G30120; PDH-E1_ALPHA, AT1G01090). Among the genes related to the Calvin-Benson-Bassham (CBB) cycle, most of the genes showed upregulation in terms of expression, and the majority of the metabolites were affected (Fig. 3C). While pentose phosphate (PenP), FBP, and T3P metabolites decreased significantly, other metabolites mostly showed an increase in their levels under the low CO_2_ condition (Fig. 3C). These results were consistent with our enrichment results, indicating *Arabidopsis* changed expression of primary carbon metabolism and adjusted their metabolite to boost their adaptability to low CO_2_ condition.

### NH_4_^+^ treatment increased the expression of genes involved in ammonia refixation

In our results, the leaf content NH_4_^+^ increased (Figure 2B) and the ammonia refixation was induced (Figure 2C) following low CO_2_ treatment. Combining assumptions proposed by (Mallmann et al., 2014), we propose that ammonia refixation induced by high rate of photorespiration, contributes to the induction of C_4_ related genes under low CO_2_ condition. The NH_4_^+^ treatment experiment was conducted and revealed that, in comparison with plants cultivated continuously under control check (CK), the treated plants with 30 mM NH_4_^+^ increased the NH_4_^+^ content by at least one-fold, and the NH_4_^+^ content increased by at least three-fold with the 60mM NH_4_^+^ treatment (Fig. 4C). RT-qPCR analysis revealed that NH_4_^+^ treatment upregulated the expression of genes involved in ammonia refixation, specifically GS2 and Fd-GOGAT as shown in Figure 4B. Interestingly, this increase was not observed when plants were treated with NH_4_CI, K_2_SO_4_, KNO_3_, or Mannitol (Li et al., 2019) (Fig. 4C). Therefore, NH_4_^+^ ion is a crucial factor that induces the expression of genes related to ammonia refixation.

**Fig. 4.**
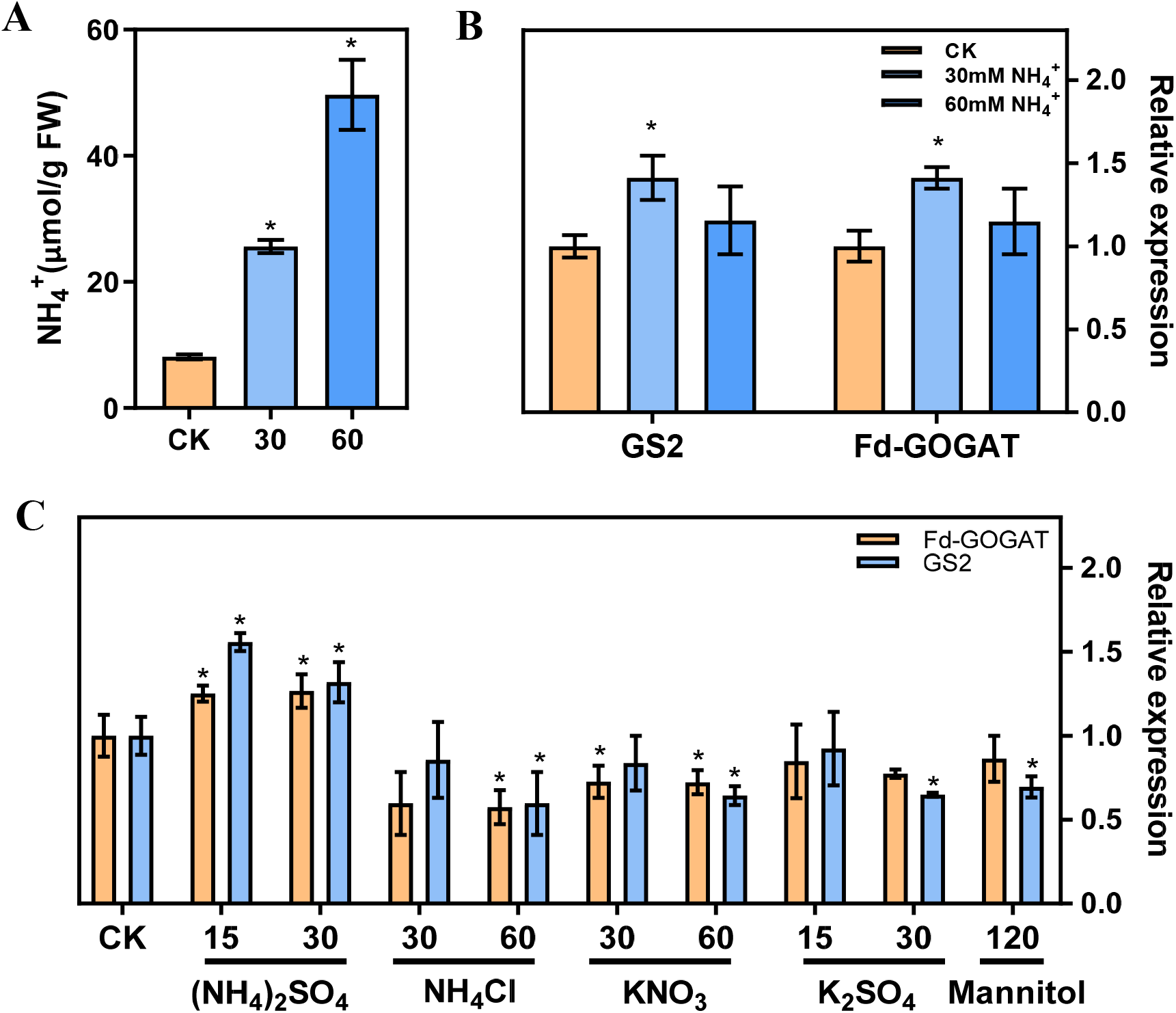
*In vitro* NH_4_^+^ treatment increased the expression of genes involved in ammonia refixation. (A) NH_4_^+^ content in the seedlings (Control check, CK; 30 mM NH_4_^+^, 30; 60 mM NH_4_^+^, 60) (replications *n*=4). (B) The gene expression of the Glutamine synthetase 2 (GS2) and ferredoxin-dependent GOGAT (Fd-GOGAT) were analyzed using RT–qPCR (replications *n*=4). (C) The effects of different ion on the gene expression of GS2 and Fd- GOGAT. Data are shown as mean ± s.d, * *p*<0.05, which were determined by two-sided Student’s *t-test*.

### Comparison of transcriptomics data between NH_4_^+^ treatment and low CO_2_ treatment

After NH_4_^+^ treatments, a total of 3920 genes were identified as differentially expressed. with 1910 genes upregulated and 2010 genes downregulated (Fig. S4A). The GO terms for upregulated genes were enriched in stress, nitrogen component metabolism, carbohydrate biosynthetic process and oxoacid metabolism (Fig. S4B). These findings are in line with earlier findings that showed that excessive NH_4_^+^ induced ammonia refixation related genes, such as glutamate dehydrogenase (Patterson et al., 2010; Daniela Ristova, 2016), and GS/GOGAT pathway (Liu and von Wiren, 2017). Likewise, the results of the enrichment analysis of KEGG show that upregulated genes were significantly enriched in carbon metabolism and 2-oxocarboxylic acid metabolism (Dataset S2), and the latter is the most elementary set of metabolites that includes pyruvate, oxaloacetate and 2-OG which enters the TCA cycle and supplement carbon source. These results indicate carbon metabolism related pathway were induced to alleviate NH_4_^+^ stress under NH_4_^+^ treatments. We compared the transcriptomic changes induced by both NH_4_^+^ and low CO_2_ treatments and found that DEGs (*P<0.05*) in both treatments showed a higher Pearson correlation coefficient than those with no significant changes to either treatment (Fig. 5A) (*P<0.001*), which suggest that the treatment of NH_4_^+^ and low CO_2_ caused highly similar changes. The Venn diagram shows that 525 genes were jointly upregulated (Fig. 5B; Dataset S3), while 464 genes were jointly downregulated (Fig. S4C) under both treatments. The jointly upregulated genes are mostly enriched in glycolysis, starch and sucrose metabolism, carbon metabolism, accounting for 36% of those differentially expressed upregulated genes (Fig. 5C). The jointly upregulated genes were also found to be enriched in biological processes related to light stimulation, water deprivation, cold, abscisic acid, and the glycolytic process (Fig. 5D). These results suggest that multiple genes involved in primary metabolism were simultaneously affected by both NH ^+^ and low CO_2_ treatments.

**Fig 5.**
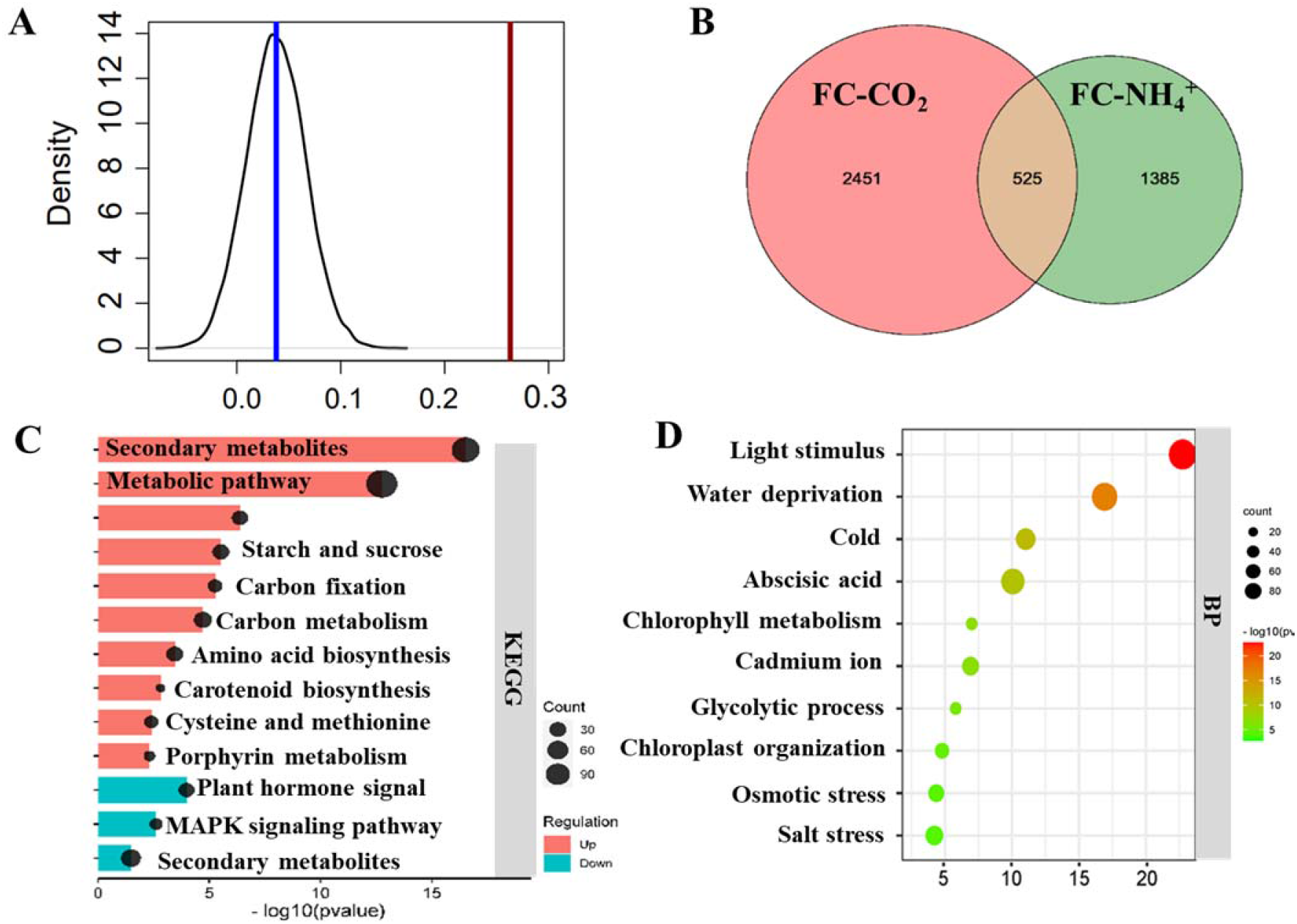
Comparison of transcriptomic profiling between NH_4_^+^ and low CO_2_ treatments. (A) Pearson correlation analysis of transcriptomics data between NH_4_^+^ and low CO_2_ treatment. The black line represents the distribution of Pearson correlations of randomly selected genes (1632) for 1000 times, and the red line represents the Pearson correlation of differentially expressed 1632 genes. A cutoff value for *adjusted P-value* < 0.05 were used to identify differentially expressed genes. (B) Venn diagrams. The diagram shows the number of genes which are jointly significantly upregulated (*adjusted P-value* < 0.05) under low CO_2_ treatment (FC-CO_2_) and NH_4_^+^ treatment (FC-NH_4_^+^). Detailed gene information can be found in Dataset S3. (C) KEGG enrichment analysis of significantly jointly upregulated genes. (D) Gene ontology (GO) Biological process (BP) enrichment analysis of significantly jointly upregulated genes. Top 10 significantly enriched results are shown here (C, D). Remaining enriched results can be found in Dataset S3.

### Influence of NH_4_^+^ treatment on the gene expression in primary metabolism

Transcriptomic data from plants treated with NH_4_^+^ showed an increase in expression levels of the GS/GOGAT pathway and NADH-dependent glutamate dehydrogenase (GDH), as well as upregulation of nitrate reductase genes (NIA1, AT1G77760; NIA2, AT1G37130) (Masclaux-Daubresse et al., 2010) (Fig. 6A), suggesting that NH_4_^+^ treatment enhances ammonia refixation metabolism. Additionally, RNA-seq data showed that almost all genes related to the photorespiratory pathway were induced, along with the expressions of genes related to the TCA cycle, Glycolysis pathway, and the CBB cycle (Fig. 6A). The transcriptomic results in primary metabolism induced by NH_4_^+^ treatment were similar to those induced by low CO_2_ treatment (Fig. S5).

**Fig. 6.**
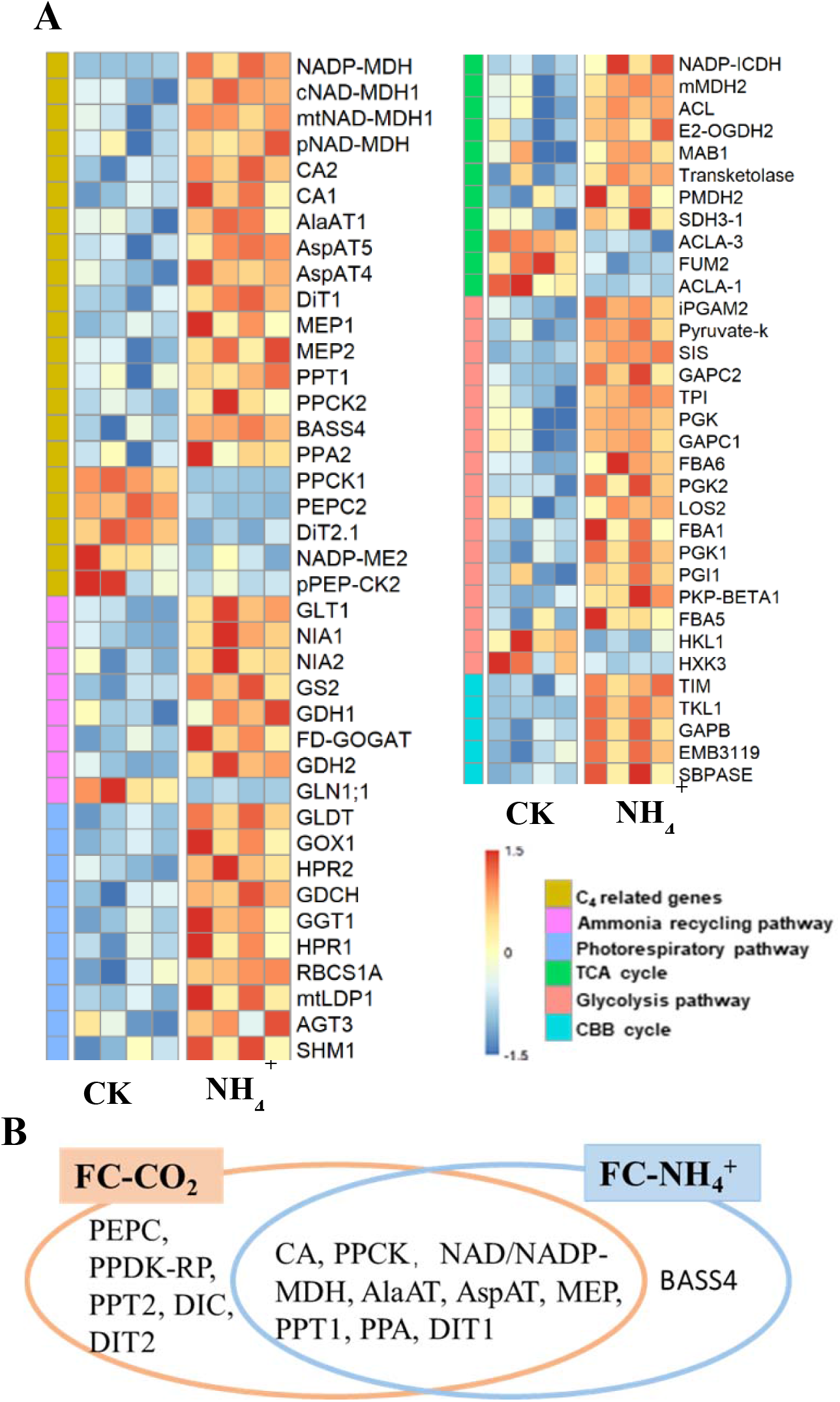
(A) Heatmap of differentially expressed genes of different metabolic pathways under NH_4_^+^ treatment (replications *n*=4). Genes from six pathways were shown as labeled in different colors indicated in the “Pathways” legend. (B) Comparative analysis of C_4_ related enzymes under low CO_2_ treatment (FC-CO_2_) and NH_4_^+^ treatment (FC-NH_4_^+^). The fisher’s exact text *P<0.05*.

Further analysis suggested that NH_4_^+^ treatment induced significantly the expression of most C_4_ related genes, including CA (CA1, AT3G01500; cytoplasmic CA2, AT5G14740), NADP-dependent malate dehydrogenase (NADP-MDH, AT5G58330), cytosolic malate dehydrogenase (cMDH1, AT1G04410), NAD-MDH1 (AT1G53240, AT3G47520), and phosphoenolpyruvate carboxylase kinase (PPCK2, AT3G04530) (Fig. 6A and Dataset S1). Furthermore, some aminotransferases, such as AspAT4, AspAT5 (AT4G31990), and AlaAT1 were up-regulated by NH_4_^+^ treatment (Fig. 6A and Dataset S1). Several transporter genes, including DiT1, PPT1, Sodium Bile acid symporter (BASS4, AT3G56160), PPA2, NHD1, and MEP, were also upregulated by NH_4_^+^ treatment (Fig. 6A and Dataset S1). However, the expression levels of PEPC, PPCK1, and pPEP-CK2 were downregulated (Fig. 6A). In addition, we also found that CA, NAD/NADP-MDH, PPCK, AlaAT, AspAT, MEP, PPT1, PPA, and DIT1 were significantly induced by both treatments (Fig 6B). Furthermore, low CO_2_ treatment induced the upregulation of most C_4_ related genes, almost including those induced by NH_4_^+^ treatment, excepte for BASS4 (Fig 6B) (Fisher’s exact text *P<0.05*). Therefore, NH_4_^+^ accumulation may be a mechanism that upregulates C_4_ genes under condition of low CO_2_ (Fig 6B).

## Discussion

### The gene expression involved in the primary metabolism is adjusted to increase the capacity of ammonia refixation under low CO_2_ condition

*Arabidopsis* grown under low CO_2_ condition showed a significant decrease in growth (Li et al., 2014) and chlorophyll content compared to the control (Fig. S1A, B). The decreased biomass can be attributed to the decreased photosynthetic CO_2_ uptake rate under low CO_2_ condition. Logically, decreased biomass production coupled with increased expression of genes related to chlorophyll synthesis should lead to increased leaf chlorophyll content. The mechanism underlying the observed decrease in chlorophyll concentration is unknown, one possibility is that under low CO_2_ certain processes damaging chlorophyll might be activated. In addition, the evolution of C_4_ photosynthesis has been hypothesized to be promoted by the decrease in atmospheric CO_2_ (Ehleringer, 1991; Helliker, 1997; Sage, 2012). The upregulation of PEPC and PEPE-K genes in response to low CO_2_ has also been observed (Li et al., 2014). Nevertheless, the contribution of low CO_2_ condition to the evolutionary assembly of the C_4_ pathway, especially for C_4_ gene expression, remains to be further systematicly studied (Li et al., 2014). In our study, we found that the expression of almost all core C_4_ cycle enzymes and transporters were upregulated under low CO_2_ treatment (Fig1). Interestingly, a similar upregulation of C_4_ like pathway genes for carbon fixation was observed in Nannochloropsis oceanica under low CO_2_ condition (Wei et al., 2019). This discovery suggests a significant role of low CO_2_ condition in the evolution of C_4_ photosynthesis and implies that the strengthening of expression of C_4_ photosynthesis genes may enhance the adaptation ability to low CO_2_ condition.

Low CO_2_ condition, like other stresses such as drought and high temperature (Carmo-Silva et al., 2008; Peterhansel and Maurino, 2011), increase photorespiration (Fig. 2A) and promote NH_4_^+^ production (Fig. 2B) (Frantz et al., 1982; Alencar et al., 2019). Photorespiration is the primary source of NH_4_^+^ in higher plants (Leegood et al., 1995; Kumagai et al., 2011). *Arabidopsis* as an NH_4_^+^ sensitive plant (Cruz et al., 2006), experiencing the detrimental effects of high levels of NH_4_^+^, i.e inhibiting plant growth and leaf chlorosis (Li et al., 2012; Li et al., 2014; Esteban et al., 2016), which are also observed under low CO_2_ condition (Fig. S1A, B). Taken together, these results suggest that *Arabidopsi*s experiences ammonia stress under low CO_2_ condition.

Previous studies have shown that excess ammonia is toxic (Hachiya et al., 2012; Esteban et al., 2016) and need to be recycled by upregulating several key enzymes (Cruz et al., 2006; Liu and von Wiren, 2017; Hachiya et al., 2021). The GS/GOGAT pathway which is involved in ammonia refixation was enhanced under low CO_2_ condition (Fig. 2*C*). The TCA cycle under illumination helps provide the carbon skeleton, in particular 2- OG, for ammonia refixation (Nunes-Nesi et al., 2010; Sweetlove et al., 2010; Tcherkez et al., 2017). The gene expression of IDH1 and ACO1 which are used for the generation of 2-OG, were also significantly upregulated in the TCA cycle (Fig. 3*A*). Although the concentration of 2-OG remained unchanged (Arrivault et al., 2009), a significant reduction in the concentrations of fumarate and succinate were observed (Fig. 3*A*), implying that there is an increased flux of 2-OG used for ammonia refixation, rather than decarboxylation in the TCA cycle. Consistent with this, studies have also shown that supplementing with succinate and 2-OG can increase ammonia recirculation and alleviate ammonia toxicity (Yuan et al., 2007; Hachiya et al., 2012).

Previous calculations suggest that the stored leaf citrate content available at the start of light period is insufficient to support the amount of 2-OG required for glutamate production (Stitt et al., 2002). The 2-OG can also be generated by reactions catalyzed by aminotransferases. In this aspect, PEPC might play a crucial role during the synthesis of additional 2-OG (Turpin, 1994), potatoes with overexpressed PEPC display increased 2- OG levels in the leaves (Thomas Rademacher1, 2002), which catalyzes the formation of OAA and participates in transamination to produce 2-OG. In line with this, the genes encoding PEPC, PEPC-K, AspAT, and MDH are also upregulated under low CO_2_ condition (Fig. 7). In addition, CAs are invovled in optimizing cytosolic PEPC activity and ensure normal growth under low CO_2_ condition (DiMario et al., 2016; Weerasooriya et al., 2022) which increased significantly under low CO_2_ condition (Fig.1). Similarly, the gene expression of PPDK-RP, and AlaAT were all upregulated to produce pyruvate and carbon skeletons (Fig.7). Besides, the related transporters such as Dit1, DIT2.1, and so on were increased to satisfy the increased needs of metabolites transport as well (Gowik et al., 2011) (Fig.7). All these show that under low CO_2_ condition, the primary metabolism can be reprogrammed to generate the required carbon skeletons for ammonia refixation.

**Fig. 7.**
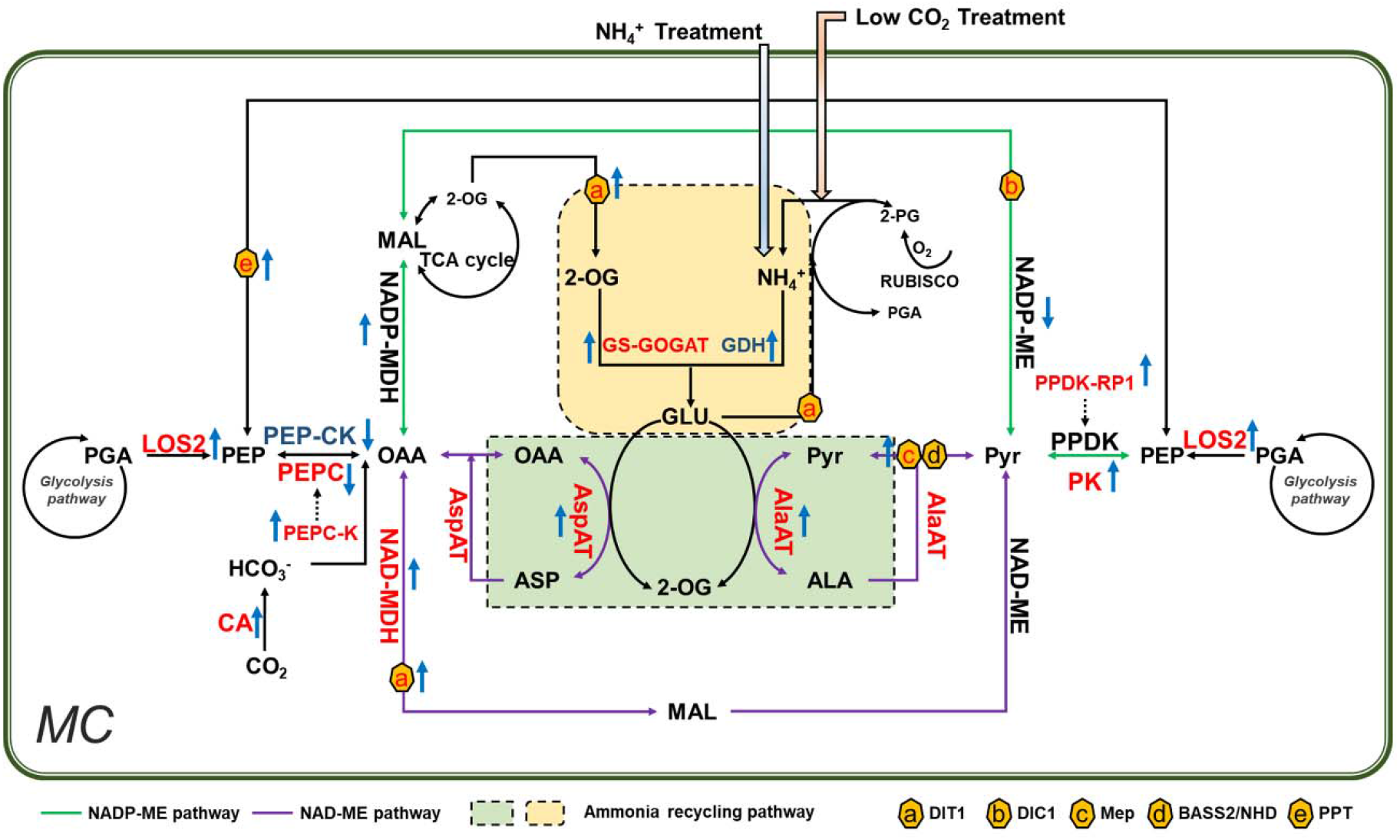
Ammonia accumulation and refixation is a primary mechanism inducing C_4_ related gene expression under low CO_2_ condition in *Arabidopsis*. Low CO_2_ treatment caused ammonia toxicity in *Arabidopsis*. Recirculation of ammonia requires reprogramming of primary metabolism reactions that involved in virtually all core C_4_ genes. *In vitro* NH_4_^+^ treatment induces a similar pattern of changes in C_4_ related genes and genes involved in primary metabolism. Therefore, we proposed that increased ammonia is a primary mechanism to induce the expression of C_4_ related gene under low CO_2_ in *Arabidopsis*. The Enzymes labeled in different colors were indicated as red/blue: upregulated/downregulated under low CO_2_. Rising/descending arrows beside the enzymes indicated upregulated/downregulated in NH_4_^+^ treatment. NADP-ME and NAD- ME pathways were represented in blue and purple lines respectively. Ammonia recycling pathways were shown in green and yellow rectangles; membrane transporters were indicated in hexagons.

### NH_4_^+^ accumulation is primary mechanism inducing change in the expression of C_4_ related genes under low CO_2_ in *Arabidopsis*

How could low CO_2_ induce the observed changes in transcriptomics? Here we discuss four possibilities. First, the phenomenon that the primary metabolism was reprogrammed under low CO_2_ condition to alleviate ammonia toxicity was observed (Fig.7), and given ammonia rebalancing between BSC and MC is theoretically possible to facilitate evolution of C_4_ photosynthesis (Mallmann et al., 2014). Therefore, it is worth investigating whether the observed changes in gene expression of primary metabolism are primarily caused by increased ammonia levels? Our result suggest that the DEGs between low CO_2_ and NH_4_^+^ treatment show correlations (Fig. 5A). Specially, the TCA cycle and related aminotransferases which provide 2-OG for ammonia refixation were significantly upregulated (Fig. 6A), as found under low CO_2_ condition (Fig. S4). Furthermore, low CO_2_ condition upregulating C_4_ related genes almost contain NH_4_^+^ treatment induced C_4_ related genes (Test of independence using Fisher test, *P<0.05*), expect BASS4 (Fig. 6B). Therefore, we proposed NH_4_^+^ accumulation and refixtion is primary mechanism inducing change in the expression of C_4_ related genes under low CO_2_ in *Arabidopsis*. However, the detailed mechanism of how ammonia induced these diverse responses is still unknown. Transcriptomic analysis reveals that among the genes activated in response to NH_4_^+^, 90% are regulated by *AMOS1* (Li et al., 2012). Interestingly, the gene expression of *AMOS1* was increased after low CO_2_ treatment (Fig. 2D), suggesting *AMOS1* might be as a potential candidate to induce C_4_ related gene expression.

Secondly, other metabolites which show dramatic changes in concentration under low CO_2_ may also induce the observed changes in transcriptomics. Flugel et al. (Flugel et al., 2017) found that 2-PG inhibits CBB cycle enzymes TPI and SBPase in *Arabidopsis* after treatment with low CO_2_. However, the role of 2-PG in regulating primary metabolism related genes in higher plants remains unknown, despite it is crucial in the regulation of *cmpA, sbtA, and ndhF3,* components of the cyanobacterial bicarbonate transport mechanism (Haimovich-Dayan et al., 2015). Thirdly, low CO_2_ causes decreased photosynthesis, which resulted in decreased biomass production (Fig. S1A) and decreased concentrations of certain metabolites, such as sucrose (Fig. 3B). In general, decreased concentration of carbohydrate enhances expression of genes for CO_2_ fixation for both C_3_ and C_4_ plants and also genes related to mobilization of photosynthesis reserve, and export process (Sheen, 1990; Cheng et al., 1992; Koch, 1996; Ruan, 2014). Given this, decreased sugar concentration might also be a potential mechanism underlining the observed reconfiguration of primary metabolism under low CO_2_ condition. In addition, according to (You et al., 2020), ABA and PEPC play crucial roles in enabling *Arabidopsis* to adapt to low CO_2_ condition. Intriguingly, there was a notable increase in the expression of ABA related metabolic processes under low CO_2_ treatment (Dataset S2) which implies that ABA signaling might have been involve in the observed changed in the transcriptomic under low CO_2_ condition as well.

### A regulatory preconditioning for the emergence of C_4_ photosynthesis from C_3_ photosynthesis

Research indicates that low CO_2_ condition were not the trigger for the evolution of C_4_ plants (Sage, 2012), which more likely served as a preconditioning that played a significant role for the C_4_ formation (Ehleringer, 1991; Osborne and Freckleton, 2009; Edwards and Smith, 2010; Sage, 2012). However, how its contribution to the formation of C_4_ plants remains unclear.

Results from this study provide a partial answer to this question. C_2_ photosynthesis, is considered as an intermediate stage during C_4_ evolution (Bauwe et al., 2010; Heckmann et al., 2013), which also causing a potential ammonia misbalance between BSC and MC (Mallmann et al., 2014). A number of putative pathways to recycle NH_4_^+^ were proposed (Mallmann et al., 2014), which usually use many, though not all, enzymes in the C_4_ cycle. In all these proposed pathways, ammonia is refixed by GS/GOGAT pathway with carbon skeleton provided by a partial C_4_ cycle. Several experimental results support this hypothesis. For example, in C_3_–C_4_ intermediate Flaveria species, the transcripts GS/GOGAT are upregulated, as also the case for a number of aminotransferases (Mallmann et al., 2014). Considering that C_4_ photosynthesis emerged from C_3_ photosynthesis in a relatively short geological time scale, i.e., only within about ∼20 million years (Christin et al., 2008), how could C_3_ plants rapidly gain such complex regulatory mechanisms regulating ammonia recycling and providing carbon skeleton when the atmospheric CO_2_ drops at Oligocene?

Our results first found and proposed that the regulatory mechanism inducing ammonia recycling pathway pre-existed in C_3_ plants. Under low CO_2_ condition or other conditions where the photorespiratory flux is high and the ammonia release flux is high, an innate regulatory mechanism to induce expression of genes involved in ammonia recycling can be activated. This is sensible since condition causing physiological low CO_2_ condition, e.g. drought, high temperature, etc. which causes stomatal closure (James R. Ehleringer, 1991; DAVIES, 2002; Sage et al., 2018), widely exist, mechanisms to enable capturing the released ammonia will confer competitive advantage for plants. This innate ammonia recycling mechanism therefore represents a regulatory preconditioning for creating different metabolic pathways to solve the challenge of ammonia misbalance in C_2_ species (Mallmann et al., 2014). Previous study found C_4_ evolution is built upon pre-existing molecular mechanisms (Aubry et al., 2011; Reyna-Llorens and Hibberd, 2017; Dickinson et al., 2020). Our study suggests that the widespread pre-existing ammonia recycling mechanism in C_3_ plants might have facilitated the polyphyletic C_4_ evolution. Therefore, we provide another evidence for the mechanism of multiple parallel evolutions of C_4_ photosynthesis and, and a better understanding of how these changes are likely to have occurred should help introduce C_4_ characteristics into C_3_.

## Supplementary data

**Fig. S1.** Growth was arrested in A. thaliana under low CO_2_ treatment

**Fig. S2.** Analysis of differentially expressed genes associated with low CO_2_ treatment

**Fig. S3.** C_4_ related genes were verified by RT-qPCR

**Fig. S4.** Identification of differentially expressed genes associated with NH_4_^+^) treatment. **Fig. S5.** The comparison of gene expression between low CO_2_ treatment (FC-CO_2_) and NH_4_^+^ (FC- NH_4_^+^) treatment in different metabolic pathways

**Table S1.** Gene-specific primers used for RT-qPC

**Datasets S1.** The differentially expressed genes (DEGs) of transcripts and primary metabolism after low CO_2_ and NH_4_^+^ treatment, and the changes in metabolic profiles under low CO_2_ treatment.

**Datasets S2.** GO and KEGG enrichment analysis after low CO_2_ and NH_4_^+^ treatment. **Datasets S3.** Overlap of DEGs after low CO_2_ and NH_4_^+^ treatment and their KEGG and GO enrichment data.

## Acknowledgments.

We thank Dr. Qingqiu Gong, Dr. Qingfeng Song, Yongyao Zhao, Xiaoxiang Ni, Yuhui Huang, Dr. Faming Chen, Dr. Qiming Tang, Dr. Genyun Chen, Jianzhao Yang for technical support and inspiring discussion during the project. We thank The Chinese Academy of Sciences (XDB27020105), Ministry of Science and Technology of China (2020YFA0907600, XDB37020104), and National Science Foundation (31870214) for financial support.

## Author Contributions

F.M., and N.U.H., performed research; Q.T., performed the metabolomics analysis; F.C., performed the RNA-seq analysis; F.M., and X.Z., wrote the paper.

## Data Availability

The RNA-seq datasets have been submitted to the NCBI Gene Expression Omnibus under accession number PRJNA842829. All other study data are included in the article and/or supporting information.

## References

1. Alencar V, Lobo AKM, Carvalho FEL, Silveira JAG (2019) High ammonium supply impairs photosynthetic efficiency in rice exposed to excess light. Photosynth Res 140: 321–335

2. Arnon DI (1949) Copper Enzymes in Isolated Chloroplasts. Polyphenoloxidase in Beta Vulgaris. Plant Physiol 24: 1–15

3. Arrivault S, Alexandre Moraes T, Obata T, Medeiros DB, Fernie AR, Boulouis A, Ludwig M, Lunn JE, Borghi GL, Schlereth A, Guenther M, Stitt M (2019) Metabolite profiles reveal interspecific variation in operation of the Calvin-Benson cycle in both C4 and C3 plants. J Exp Bot 70: 1843–1858

4. Arrivault S, Guenther M, Ivakov A, Feil R, Vosloh D, van Dongen JT, Sulpice R, Stitt M (2009) Use of reverse-phase liquid chromatography, linked to tandem mass spectrometry, to profile the Calvin cycle and other metabolic intermediates in Arabidopsis rosettes at different carbon dioxide concentrations. Plant J 59: 826–839

5. Aubry S, Brown NJ, Hibberd JM (2011) The role of proteins in C(3) plants prior to their recruitment into the C(4) pathway. J Exp Bot 62: 3049–3059

6. Bauwe H, Hagemann M, Fernie AR (2010) Photorespiration: players, partners and origin. Trends Plant Sci 15: 330–336

7. Carmo-Silva AE, Powers SJ, Keys AJ, Arrabaca MC, Parry MA (2008) Photorespiration in C4 grasses remains slow under drought conditions. Plant Cell Environ 31: 925–940

8. Chen JH, Chen ST, He NY, Wang QL, Zhao Y, Gao W, Guo FQ (2020) Nuclear-encoded synthesis of the D1 subunit of photosystem II increases photosynthetic efficiency and crop yield. Nat Plants 6: 570–580

9. Cheng CL, Acedo GN, Cristinsin M, Conkling MA (1992) Sucrose mimics the light induction of Arabidopsis nitrate reductase gene transcription. Proc Natl Acad Sci U S A 89: 1861–1864

10. Christin PA, Besnard G, Samaritani E, Duvall MR, Hodkinson TR, Savolainen V, Salamin N (2008) Oligocene CO2 decline promoted C4 photosynthesis in grasses. Curr Biol 18: 37–43

11. Cruz C, Bio AF, Dominguez-Valdivia MD, Aparicio-Tejo PM, Lamsfus C, Martins-Loucao MA (2006) How does glutamine synthetase activity determine plant tolerance to ammonium? Planta 223: 1068–1080

12. Daniela Ristova, 2 Clément Carré,3,4 Marjorie Pervent,3 Anna Medici,3 Grace Jaeyoon Kim,1 Domenica Scalia,1 Sandrine Ruffel,3 Kenneth D. Birnbaum,1 Benoît Lacombe,3 Wolfgang Busch,2 Gloria M. Coruzzi,1 Gabriel Krouk3 (2016) Combinatorial interaction network of transcriptomic and phenotypic responses to nitrogen and hormones in the Arabidopsis thaliana root.

13. Davies SWWJ (2002) ABA-based chemical signalling: the co-ordination of responses to stress in plants.

14. Dickinson PJ, Knerova J, Szecowka M, Stevenson SR, Burgess SJ, Mulvey H, Bagman AM, Gaudinier A, Brady SM, Hibberd JM (2020) A bipartite transcription factor module controlling expression in the bundle sheath of Arabidopsis thaliana. Nat Plants 6: 1468–1479

15. DiMario RJ, Quebedeaux JC, Longstreth DJ, Dassanayake M, Hartman MM, Moroney JV (2016) The Cytoplasmic Carbonic Anhydrases betaCA2 and betaCA4 Are Required for Optimal Plant Growth at Low CO2. Plant Physiol 171: 280–293

16. Dobin A, Davis CA, Schlesinger F, Drenkow J, Zaleski C, Jha S, Batut P, Chaisson M, Gingeras TR (2013) STAR: ultrafast universal RNA-seq aligner. Bioinformatics 29: 15–21

17. Edwards EJ, Smith SA (2010) Phylogenetic analyses reveal the shady history of C4 grasses. Proc Natl Acad Sci U S A 107: 2532–2537

18. Ehleringer JR, Cerling TE, Helliker BR (1997) C4 photosynthesis, atmospheric CO2, and climate. Oecologia 112: 285–299

19. Ehleringer JR, Sage RF, Flanagan LB, Pearcy RW (1991) Climate Change and the Evolution of C4 Photosynthesis. Trends in Ecology & Evolution 6: 95–99

20. Ehleringer JR, Sage, R. F., Flanagan, L. B., and Pearcy, R. W (1991) Climate change and the evolution of C4 photosynthesis. .

21. Esteban R, Ariz I, Cruz C, Moran JF (2016) Review: Mechanisms of ammonium toxicity and the quest for tolerance. Plant Sci 248: 92–101

22. Flugel F, Timm S, Arrivault S, Florian A, Stitt M, Fernie AR, Bauwe H (2017) The Photorespiratory Metabolite 2-Phosphoglycolate Regulates Photosynthesis and Starch Accumulation in Arabidopsis. Plant Cell 29: 2537–2551

23. Frantz TA, Peterson DM, Durbin RD (1982) Sources of ammonium in oat leaves treated with tabtoxin or methionine sulfoximine. Plant Physiol 69: 345–348

24. Gowik U, Brautigam A, Weber KL, Weber AP, Westhoff P (2011) Evolution of C4 photosynthesis in the genus Flaveria: how many and which genes does it take to make C4? Plant Cell 23: 2087–2105

25. Hachiya T, Inaba J, Wakazaki M, Sato M, Toyooka K, Miyagi A, Kawai-Yamada M, Sugiura D, Nakagawa T, Kiba T, Gojon A, Sakakibara H (2021) Excessive ammonium assimilation by plastidic glutamine synthetase causes ammonium toxicity in Arabidopsis thaliana. Nat Commun 12: 4944

26. Hachiya T, Watanabe CK, Fujimoto M, Ishikawa T, Takahara K, Kawai-Yamada M, Uchimiya H, Uesono Y, Terashima I, Noguchi K (2012) Nitrate addition alleviates ammonium toxicity without lessening ammonium accumulation, organic acid depletion and inorganic cation depletion in Arabidopsis thaliana shoots. Plant Cell Physiol 53: 577–591

27. Hachiya T, Watanabe CK, Fujimoto M, Ishikawa T, Takahara K, Kawai-Yamada M, Uchimiya H, Uesono Y, Terashima I, Noguchi K (2012) Nitrate Addition Alleviates Ammonium Toxicity Without Lessening Ammonium Accumulation, Organic Acid Depletion and Inorganic Cation Depletion in Arabidopsis thaliana Shoots. Plant and Cell Physiology 53: 577–591

28. Haimovich-Dayan M, Lieman-Hurwitz J, Orf I, Hagemann M, Kaplan A (2015) Does 2- phosphoglycolate serve as an internal signal molecule of inorganic carbon deprivation in the cyanobacterium Synechocystis sp. PCC 6803? Environ Microbiol 17: 1794-1804

29. Heckmann D, Schulze S, Denton A, Gowik U, Westhoff P, Weber AP, Lercher MJ (2013) Predicting C4 photosynthesis evolution: modular, individually adaptive steps on a Mount Fuji fitness landscape. Cell 153: 1579–1588

30. Helliker JRETECBR (1997) C4 photosynthesis, atmospheric CO2 and climate.

31. James R. Ehleringer RFS, Lawrence B. Flanagan and Robert W. Pearcy (1991) Climate Change and the Evolution of C4 Photosynthsis.

32. Keerberg O, Parnik T, Ivanova H, Bassuner B, Bauwe H (2014) C2 photosynthesis generates about 3-fold elevated leaf CO2 levels in the C3-C4 intermediate species Flaveria pubescens. J Exp Bot 65: 3649–3656

33. Keys AJ (2006) The re-assimilation of ammonia produced by photorespiration and the nitrogen economy of C3 higher plants. Photosynth Res 87: 165–175

34. Keys AJ, Bird IF, Cornelius MJ, Lea PJ, Wallsgrove RM, Miflin BJ (1978) Photorespiratory Nitrogen Cycle. Nature 275: 741–743

35. Kinoshita H, Nagasaki J, Yoshikawa N, Yamamoto A, Takito S, Kawasaki M, Sugiyama T, Miyake H, Weber APM, Taniguchi M (2011) The chloroplastic 2-oxoglutarate/malate transporter has dual function as the malate valve and in carbon/nitrogen metabolism. Plant J 65: 15–26

36. Koch KE (1996) Carbohydrate-Modulated Gene Expression in Plants. Annu Rev Plant Physiol Plant Mol Biol 47: 509–540

37. Krogmann DW, Jagendorf AT, Avron M (1959) Uncouplers of Spinach Chloroplast Photosynthetic Phosphorylation. Plant Physiol 34: 272–277

38. Kumagai E, Araki T, Hamaoka N, Ueno O (2011) Ammonia emission from rice leaves in relation to photorespiration and genotypic differences in glutamine synthetase activity. Ann Bot 108: 1381–1386

39. Lam HM, Coschigano KT, Oliveira IC, Melo-Oliveira R, Coruzzi GM (1996) The Molecular-Genetics of Nitrogen Assimilation into Amino Acids in Higher Plants. Annu Rev Plant Physiol Plant Mol Biol 47: 569–593

40. Lancien M, Gadal P, Hodges M (2000) Enzyme redundancy and the importance of 2- oxoglutarate in higher plant ammonium assimilation. Plant Physiol 123: 817–824

41. Leegood RC, Lea PJ, Adcock MD, Häusler RE (1995) The regulation and control of photorespiration. Journal of Experimental Botany 46: 1397–1414

42. Li B, Li G, Kronzucker HJ, Baluska F, Shi W (2014) Ammonium stress in Arabidopsis: signaling, genetic loci, and physiological targets. Trends Plant Sci 19: 107–114

43. Li B, Li Q, Xiong L, Kronzucker HJ, Kramer U, Shi W (2012) Arabidopsis plastid AMOS1/EGY1 integrates abscisic acid signaling to regulate global gene expression response to ammonium stress. Plant Physiol 160: 2040–2051

44. Li G, Li B, Dong G, Feng X, Kronzucker HJ, Shi W (2013) Ammonium-induced shoot ethylene production is associated with the inhibition of lateral root formation in Arabidopsis. J Exp Bot 64: 1413–1425

45. Li G, Zhang L, Wang M, Di D, Kronzucker HJ, Shi W (2019) The Arabidopsis AMOT1/EIN3 gene plays an important role in the amelioration of ammonium toxicity. J Exp Bot 70: 1375–1388

46. Li G, Zhang L, Wang M, Di D, Kronzucker HJ, Shi W (2019) The Arabidopsis AMOT1/EIN3 gene plays an important role in the amelioration of ammonium toxicity. J Exp Bot

47. Li Y, Xu J, Haq NU, Zhang H, Zhu XG (2014) Was low CO2 a driving force of C4 evolution: Arabidopsis responses to long-term low CO2 stress. J Exp Bot 65: 3657–3667

48. Liu Y, von Wiren N (2017) Ammonium as a signal for physiological and morphological responses in plants. J Exp Bot 68: 2581–2592

49. Love MI, Huber W, Anders S (2014) Moderated estimation of fold change and dispersion for RNA-seq data with DESeq2. Genome Biol 15: 550

50. Mallmann J, Heckmann D, Brautigam A, Lercher MJ, Weber AP, Westhoff P, Gowik U (2014) The role of photorespiration during the evolution of C4 photosynthesis in the genus Flaveria. Elife 3: e02478

51. Masclaux-Daubresse C, Daniel-Vedele F, Dechorgnat J, Chardon F, Gaufichon L, Suzuki A (2010) Nitrogen uptake, assimilation and remobilization in plants: challenges for sustainable and productive agriculture. Ann Bot 105: 1141–1157

52. Nagatani H, Shimizu M, Valentine RC (1971) The mechanism of ammonia assimilation in nitrogen fixing Bacteria. Arch Mikrobiol 79: 164–175

53. Nunes-Nesi A, Fernie AR, Stitt M (2010) Metabolic and signaling aspects underpinning the regulation of plant carbon nitrogen interactions. Mol Plant 3: 973–996

54. Osborne CP, Freckleton RP (2009) Ecological selection pressures for C4 photosynthesis in the grasses. Proc Biol Sci 276: 1753–1760

55. Patterson K, Cakmak T, Cooper A, Lager I, Rasmusson AG, Escobar MA (2010) Distinct signalling pathways and transcriptome response signatures differentiate ammonium- and nitrate-supplied plants. Plant Cell Environ 33: 1486–1501

56. Peterhansel C, Maurino VG (2011) Photorespiration redesigned. Plant Physiol 155: 49–55

57. Rawsthorne S, Hylton CM, Smith AM, Woolhouse HW (1988) Photorespiratory Metabolism and Immunogold Localization of Photorespiratory Enzymes in Leaves of C-3 and C-3-C-4 Intermediate Species of Moricandia. Planta 173: 298–308

58. Reyna-Llorens I, Hibberd JM (2017) Recruitment of pre-existing networks during the evolution of C4 photosynthesis. Philos Trans R Soc Lond B Biol Sci 372

59. Ruan YL (2014) Sucrose metabolism: gateway to diverse carbon use and sugar signaling. Annu Rev Plant Biol 65: 33–67

60. Sage ASRRF (2011) C4 Photosynthesis and Related CO2 Concentrating Mechanisms.

61. Sage RF (2001) Environmental and Evolutionary Preconditionsfor the Origin and Diversification of the C4PhotosyntheticSyndrome. Plant Biology 3: 202–213

62. Sage RF, Christin PA, Edwards EJ (2011) The C(4) plant lineages of planet Earth. J Exp Bot 62: 3155–3169

63. Sage RF, Monson RK, Ehleringer JR, Adachi S, Pearcy RW (2018) Some like it hot: the physiological ecology of C4 plant evolution. Oecologia 187: 941–966

64. Sage RF, Sage TL, Pearcy RW, Borsch T (2007) The taxonomic distribution of C4 photosynthesis in Amaranthaceae sensu stricto. Am J Bot 94: 1992–2003

65. Sage RFS, T. L.Kocacinar, F. (2012) Photorespiration and the evolution of C4 photosynthesis. Annu Rev Plant Biol 63: 19–47

66. Sarasketa A, Gonzalez-Moro MB, Gonzalez-Murua C, Marino D (2014) Exploring ammonium tolerance in a large panel of Arabidopsis thaliana natural accessions. J Exp Bot 65: 6023–6033

67. Schluter U, Brautigam A, Gowik U, Melzer M, Christin PA, Kurz S, Mettler-Altmann T, Weber AP (2017) Photosynthesis in C3-C4 intermediate Moricandia species. J Exp Bot 68: 191–206

68. Schluter U, Weber APM (2020) Regulation and Evolution of C4 Photosynthesis. Annu Rev Plant Biol

69. Sheen J (1990) Metabolic repression of transcription in higher plants. Plant Cell 2: 1027–1038

70. SolÓRzano L (1969) DETERMINATION OF AMMONIA IN NATURAL WATERS BY THE PHENOLHYPOCHLORITE METHOD 1 1 This research was fully supported by U.S. Atomic Energy Commission Contract No. ATS (11-1) GEN 10, P.A. 20. Limnology and Oceanography 14: 799–801

71. Still CJ, Berry JA, Collatz GJ, DeFries RS (2003) Global distribution of C3and C4vegetation: Carbon cycle implications. Global Biogeochemical Cycles 17: 6–1-6-14

72. Stitt M, Muller C, Matt P, Gibon Y, Carillo P, Morcuende R, Scheible WR, Krapp A (2002) Steps towards an integrated view of nitrogen metabolism. J Exp Bot 53: 959–970

73. Sweetlove LJ, Beard KF, Nunes-Nesi A, Fernie AR, Ratcliffe RG (2010) Not just a circle: flux modes in the plant TCA cycle. Trends Plant Sci 15: 462–470

74. Tcherkez G, Gauthier P, Buckley TN, Busch FA, Barbour MM, Bruhn D, Heskel MA, Gong XY, Crous KY, Griffin K, Way D, Turnbull M, Adams MA, Atkin OK, Farquhar GD, Cornic G (2017) Leaf day respiration: low CO2 flux but high significance for metabolism and carbon balance. New Phytol 216: 986–1001

75. Thomas Rademacher1 y, Rainer E. Häusler2, Heinz-Josef Hirsch1, Li Zhang1, Volker Lipka1,z, Dagmar Weier1,Fritz Kreuzaler1 and Christoph Peterhänsel1, (2002) An engineered phosphoenolpyruvate carboxylase redirects carbon and nitrogen flow in transgenic potato plants.

76. Turpin CHaDH (1994) INTEGRAT ION OF CARBON AND NI TROGEN ME TABOL ISM IN PLANT AND ALGAL CELLS.

77. Vogan PJ, Frohlich MW, Sage RF (2007) The functional significance of C3-C4 intermediate traits in Heliotropium L. (Boraginaceae): gas exchange perspectives. Plant Cell Environ 30: 1337–1345

78. Wang L, Czedik-Eysenberg A, Mertz RA, Si Y, Tohge T, Nunes-Nesi A, Arrivault S, Dedow LK, Bryant DW, Zhou W, Xu J, Weissmann S, Studer A, Li P, Zhang C, LaRue T, Shao Y, Ding Z, Sun Q, Patel RV, Turgeon R, Zhu X, Provart NJ, Mockler TC, Fernie AR, Stitt M, Liu P, Brutnell TP (2014) Comparative analyses of C(4) and C(3) photosynthesis in developing leaves of maize and rice. Nat Biotechnol 32: 1158–1165

79. Weerasooriya HN, DiMario RJ, Rosati VC, Rai AK, LaPlace LM, Filloon VD, Longstreth DJ, Moroney JV (2022) Arabidopsis plastid carbonic anhydrase betaCA5 is important for normal plant growth. Plant Physiol 190: 2173–2186

80. Wei L, El Hajjami M, Shen C, You W, Lu Y, Li J, Jing X, Hu Q, Zhou W, Poetsch A, Xu J (2019) Transcriptomic and proteomic responses to very low CO2 suggest multiple carbon concentrating mechanisms in Nannochloropsis oceanica. Biotechnol Biofuels 12: 168

81. Westhoff P, Gowik U (2004) Evolution of c4 phosphoenolpyruvate carboxylase. Genes and proteins: a case study with the genus Flaveria. Ann Bot 93: 13–23

82. Wingler A, Quick WP, Bungard RA, Bailey KJ, Lea PJ, Leegood RC (1999) The role of photorespiration during drought stress: an analysis utilizing barley mutants with reduced activities of photorespiratory enzymes. Plant, Cell and Environment 22: 361–373

83. You L, Zhang J, Li L, Xiao C, Feng X, Chen S, Guo L, Hu H (2020) Involvement of abscisic acid, ABI5, and PPC2 in plant acclimation to low CO2. J Exp Bot

84. Yuan L, Loque D, Kojima S, Rauch S, Ishiyama K, Inoue E, Takahashi H, von Wiren N (2007) The organization of high-affinity ammonium uptake in Arabidopsis roots depends on the spatial arrangement and biochemical properties of AMT1-type transporters. Plant Cell 19: 2636–2652

